# Molecular Dynamics Study on the Effects of Charged Amino Acid Distribution Under low pH Condition to the Unfolding of Hen Egg White Lysozyme and Formation of Beta Strands

**DOI:** 10.1101/2021.03.25.436950

**Authors:** Husnul Fuad Zein, Ibra Alam, Piyapong Asanithi, Thana Sutthibutpong

## Abstract

Aggregation of unfolded or misfolded proteins into amyloid fibrils can cause various diseases in humans. However, the fibrils synthesized in vitro can be developed toward useful biomaterials under some physicochemical conditions. In this study, atomistic molecular dynamics simulations were performed to address the mechanism of beta-sheet formation of the unfolded hen egg white lysozyme (HEWL) under a high temperature and low pH. Simulations of the protonated HEWL at pH 2 and the non-protonated HEWL at pH 7 were performed at the highly elevated temperature of 450 K to accelerate the unfolding, followed by the 333 K temperature in some previous in vitro studies. The simulations showed that HEWL unfolded faster and refolded into structures with higher beta-strand contents at pH 2. The mechanism of beta-strand formation at the earlier stage of amyloidosis was addressed in terms of the radial distribution of amino acids, affected by the high protonation level at a low pH.

## 1. Introduction

Lysozymes are a family of globular enzymes in the immune systems of animals. A lysozyme molecule is a single polypeptide of around 130 residues that can partially hydrolyze the peptidoglycans of gram-positive bacterial cell walls [1,2]. Lysozyme is one of the protein types that are associated with the formation of amyloid fibrils under some specific conditions. The failure of specific peptides or proteins to fold or to remain correctly folded triggered non-functional protein aggregation and amyloidosis [3]. Amyloid aggregation is a hallmark of several degenerative diseases, such as Alzheimer’s disease, Parkinson’s disease, type II diabetes, Creutzfeldt Jakob, Huntington’s, amyotrophic lateral sclerosis (ALS) [4,5]. However, it was suggested that nontoxic forms of amyloid fibrils could be utilized in some applications, such as bioengineering, biosensor, drug delivery, regenerative medicine, cell-encapsulating materials, tissue engineering, molecular and electronic devices, etc. [6–17].

Amyloid fibril formation *in vitro* can be controlled under some suitable environments by several factors that either stimulate or inhibit aggregation [18]. Hen egg-white lysozyme (HEWL) is one the most commonly used proteins in protein aggregation research due to its well-characterized structure and low cost. Unfolding and refolding of HEWL during amyloidosis could be accelerated under a high temperature and acidic condition. A previous study showed that amyloid fibrils were formed when incubating HEWL under the temperature of 65 °C and pH 2 for 196 hours [1].

Conformational changes of the unfolded, refolded, and aggregated proteins can be monitored by using several methods, such as atomic force microscopy (AFM), Raman spectroscopy, and electrochemical impedance spectroscopy (EIS) [19–21]. Under an elevated temperature and acidic condition, AFM could provide images of lysozyme aggregation at various incubation times by depicting the time-dependent sizes of self-assembled spheroidal oligomers. The changing of both the secondary and tertiary structures of the aqueous lysozyme could also be observed by the shifting of Raman spectra in the range of 650 – 1875 cm^-1^ from its native state [1]. Signals from EIS spectroscopy are highly sensitive to the changing of both secondary and tertiary structures of the protein due to their unique charge transfer resistances (Rct) [21]. These methods, along with dynamic light scattering, size exclusion chromatography, transmission electron microscopy (TEM), Fourier transform-infrared (FT-IR) spectroscopy, the surface plasmon resonance (SPR) [1,22,23], have their own advantages but still lack understanding in the molecular details of protein structural changes.

Computer simulations have become an alternative tool to provide more insight into the structures and dynamics of proteins at different states under different conditions in atomistic details. The accuracy of atomistic molecular dynamics (MD) simulations to predict the molecular behavior of proteins under extreme conditions [24,25] has been improved with the continuing development of molecular mechanics forcefield parameters [26,27]. Therefore, a number of computational studies were carried out to observe the effects of temperature, solvents, and external perturbation on the unfolding and aggregation of both human lysozymes and HEWL [25,28–34]. From MD simulations performed by Moraitakis *et al*., replacing an aspartic acid residue with a histidine caused human lysozyme to unfold significantly faster under high temperature and could reproduce an experimental result. The mutation was proposed as a possible seed for amyloidosis [25]. Jafari *et al*. demonstrated in molecular detail that the lysozyme unfolds better in high concentrations of the sodium dodecyl sulfate (SDS) surfactant at 370 K, higher than the thermal denaturation midpoint temperature (Tm) [28]. Jiang *et al*. reported that high electric fields could enhance the possibility of protein unfolding due to the heterogeneous nature of charge distribution within proteins [29].

In our study, the effects of pH on the propensity of refolding and formation of beta-sheets that might lead to amyloidosis were addressed in atomistic details as the low pH condition was previously reported to facilitate amyloidosis. Additionally, the altered electrostatic properties due to the addition of positive charges during the protonation of some amino acids should affect denaturation and beta-strand formation. A series of MD simulations were performed under high temperature to accelerated unfolding processes, followed by simulations under an optimum temperature for amyloidosis reported in previous studies. Then, the conformational analysis was performed to characterize the simulated protein structures and provide the detailed mechanisms of beta-sheet formation under low pH and molecular insight of accelerating amyloidosis *in vitro* for further applications.

## 2. Methodology

GROMACS 2019.6 simulation package was used to carry out the Molecular Dynamics simulation. The initial atomistic structure of hen-egg white lysozyme was obtained from Protein Data Bank (PDB ID: 1AKI). In order to emulate pH conditions, protein coordinates were input to the PropKa software [35,36] to estimate pKa values for aspartic acid, glutamic acid, and histidine residues. If the specified pH was lower than the pKa of a residue, the protonation state was assigned to that residue. At the pH 2 condition, all histidines, glutamic acids, and aspartic acids were fully protonated. Meanwhile, at pH 7, all the aforementioned amino acids were deprotonated. The lysozyme structures at both pH conditions were solvated by using the SPC216 water model within simulation boxes of size 6.9 x 6.9 x 6.9 nm^3^, which was large enough to cover a whole protein molecule with 1 nm buffer distance. As HEWL in both protonated and deprotonated forms were positively charged, all systems were neutralized by adding Cl-counter-ions. For each simulation, the energy minimization was performed with the steepest descent algorithm for the maximum number of steps of 50000. The short-range electrostatic cutoff distance and the short-range Van der Waals cutoff distance were 1.0 nm. PME treatment was used to calculate long-range electrostatic interactions. Then, an NPT equilibration stage was performed at T = 300 K and P = 1 bar for 200 ns, which was long enough to accommodate the conformation changes from protonation states introduced at pH 2. After that, three replicas of MD simulations at 450 K were performed for 100 ns to accelerate unfolding processes of both protonated (pH 2) and deprotonated (pH 7) structures. Then, all six simulations were continued at 333 K for 200 ns to emulate the *in vitro* condition that beta-strand formation was observed. The summary of all MD simulations in this study is shown in Table 1.

**Table 1.** All simulations in this study.

| pH | protein | replica | Temperature (K) | Duration (ns) | Dimension (nm<sup>3</sup>) |
| --- | --- | --- | --- | --- | --- |
| 2 | Protonated-HEWL | R1 | 450 K | 100 ns | 6.9 x 6.9 x 6.9 nm<sup>3</sup> |
|  |  |  | 333 K | 200 ns |  |
|  | Protonated-HEWL | R2 | 450 K | 100 ns | 6.9 x 6.9 x 6.9 nm<sup>3</sup> |
|  |  |  | 333 K | 200 ns |  |
|  | Protonated-HEWL | R3 | 450 K | 100 ns | 6.9 x 6.9 x 6.9 nm<sup>3</sup> |
|  |  |  | 333 K | 200 ns |  |
| 7 | HEWL | R1 | 450 K | 100 ns | 6.9 x 6.9 x 6.9 nm<sup>3</sup> |
|  |  |  | 333 K | 200 ns |  |
|  | HEWL | R2 | 450 K | 100 ns | 6.9 x 6.9 x 6.9 nm<sup>3</sup> |
|  |  |  | 333 K | 200 ns |  |
|  | HEWL | R3 | 450 K | 100 ns | 6.9 x 6.9 x 6.9 nm<sup>3</sup> |
|  |  |  | 333 K | 200 ns |  |

All MD trajectory replicas at both pH conditions were then analyzed by the root mean square deviation (RMSD) calculations to quantify the level of protein unfolding and refolding from the global conformational changes compared with the reference PDB structure. The time-dependent information of the secondary structure content of lysozymes at both pH 2 and pH 7 were analyzedanalyzed by the DSSP algorithm [37], which identified the type of secondary structure for all regions within the protein. During the unfolding process at 450 K and the refolding processes at 333 K, the states of backbone torsions and alpha-beta transitions were monitored by Ramachandran plots for all simulation replicas at both pH conditions. The radius of gyration (Rg) was calculated as functions of time for different amino acid groups of the HEWL to monitor the distribution of charged amino acids and hydrophobic amino acids during the protein unfolding and refolding processes.

## 3. Results

To investigate the conformational change of lysozyme under different pH, root mean square deviation (RMSD) of HEWL was determined from MD trajectories. The time evolution of RMSD of all MD replicas at pH 2 and pH 7 in Figure 1 was used to describe conformational stability at two different temperature levels. The first 100 ns of each RMSD calculation represented the simulation of lysozyme at 450 K, while the last 200 ns represented the simulation at 333 K. The data was displayed in different colors - black (R0), red (R1), and green (R2) - that refer three different repeats of the simulations. In pH 2 condition at 450 K (Figure 1a), the RMSD values of lysozyme in all repeats increased rapidly within the first 30 ns to 1.3 nm, suggesting that the protein started to unfold from the beginning of the simulation and shortly became stable at 30 ns. The time and ensemble average of the equilibrated RMSD at 450 K and pH 2 was 1.35 ± 0.15 nm. RMSD values of the simulated HEWL were decreased and became more stable after the systems were cooled down to 333 K, as the averaged RMSD was found at 1.30 ± 0.10 nm. At this stage, the slight drop of RMSD values denoted the refolding of proteins. For simulations at pH 7 and 450 K (Figure 1b), the RMSD increased to around 1.30 ± 0.15 nm before reaching an equilibrium at around 50 ns, 20 ns slower than those at pH 2. The RMSD results at a very high temperature of 450 K implied that the lysozyme structure unfolded faster at the pH 2 condition compared to the lysozyme at the pH 7 condition. No significant drop of RMSD was seen when switching the temperature to 333 K, and hence no protein refolding.

**Figure 1.**
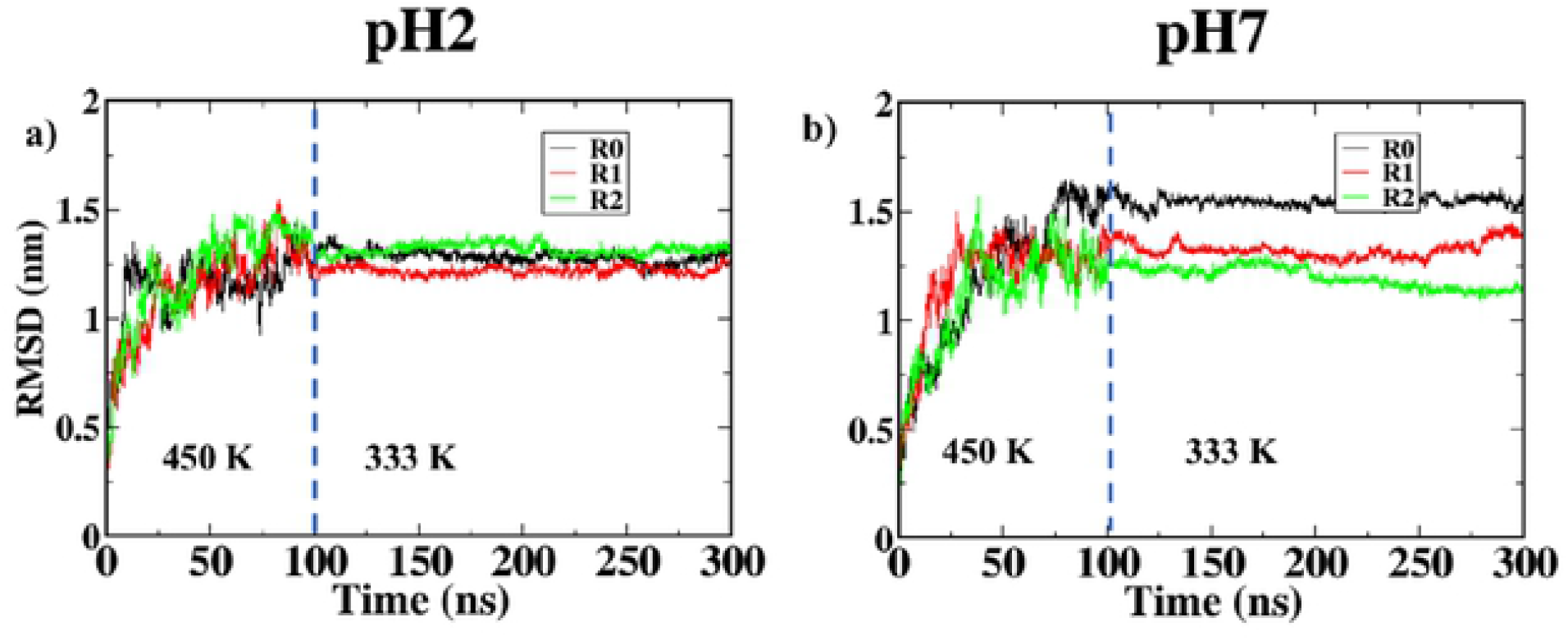
RMSD calculated as functions of time for three replicas of HEWL simulations at (a) pH 2 and (b) pH 7. Vertical dashed lines represent the time t = 100 ns where the temperature was switched from 450 K to 333 K.

Time evolution of the conformational features of HEWL at pH 2 under a very high temperature 450 K for 100 ns was monitored to observe protein unfolding and at the temperature 333 K for 200 ns to observe refolding. At pH 2 (Figure 2), most of the alpha-helical structures were lost, and the formation of beta-strands was seen towards the ends of all simulation replicas. However, at pH 7 (Figure 3), most of the secondary structures of HEWL were transformed into random coils, and only a few alpha-helical structures were observed.

**Figure 2.**
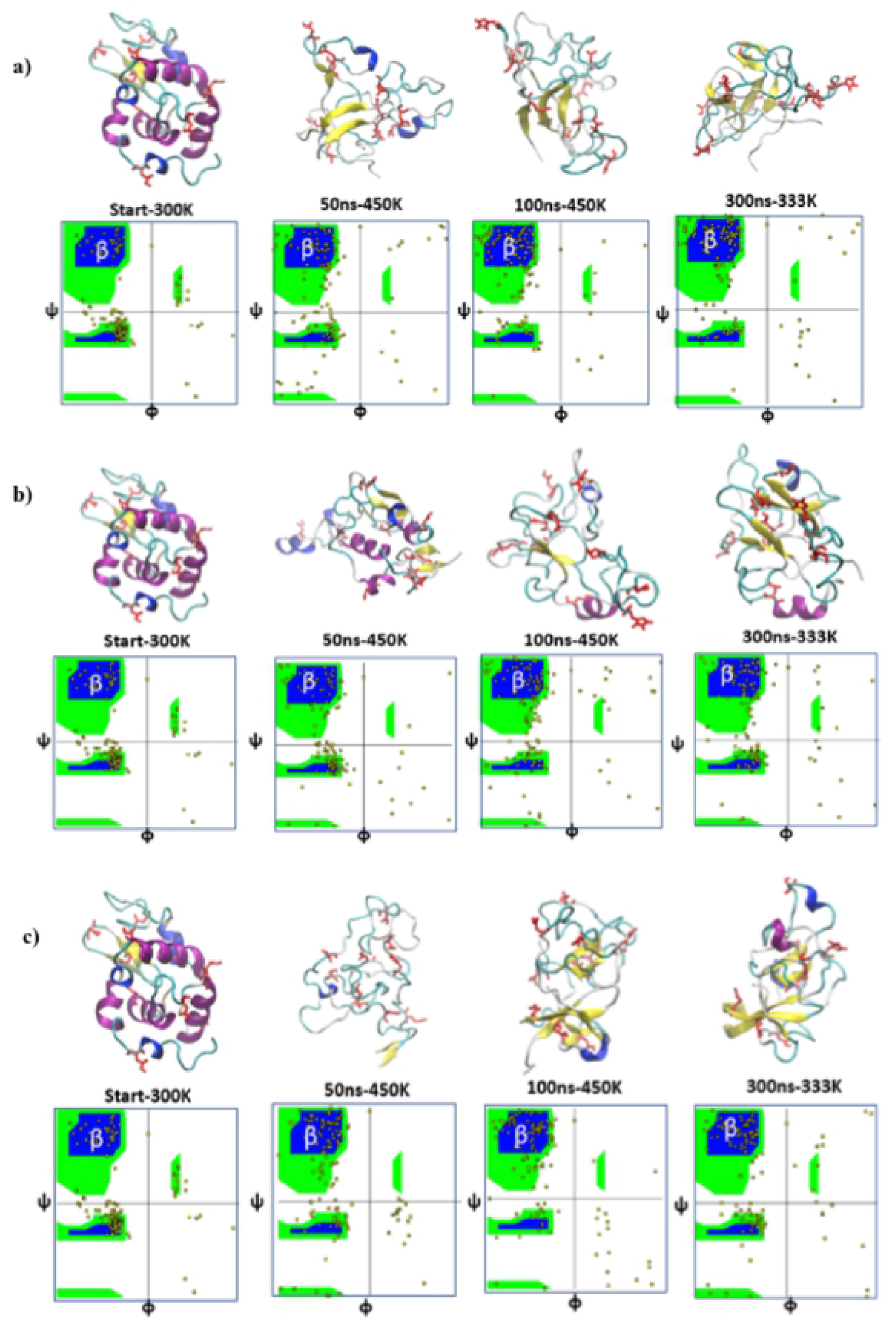
Conformational snapshots of HEWL at pH2 (top) and ramachandran plots (bottom) for the replicas (a) R0, (b) R1, and (c) R2 taken from at the start, after 50 ns of simulations at 450 K, after 100 ns of simulations at 450 K, and after 300 ns of simulations at 333 K.

**Figure 3.**
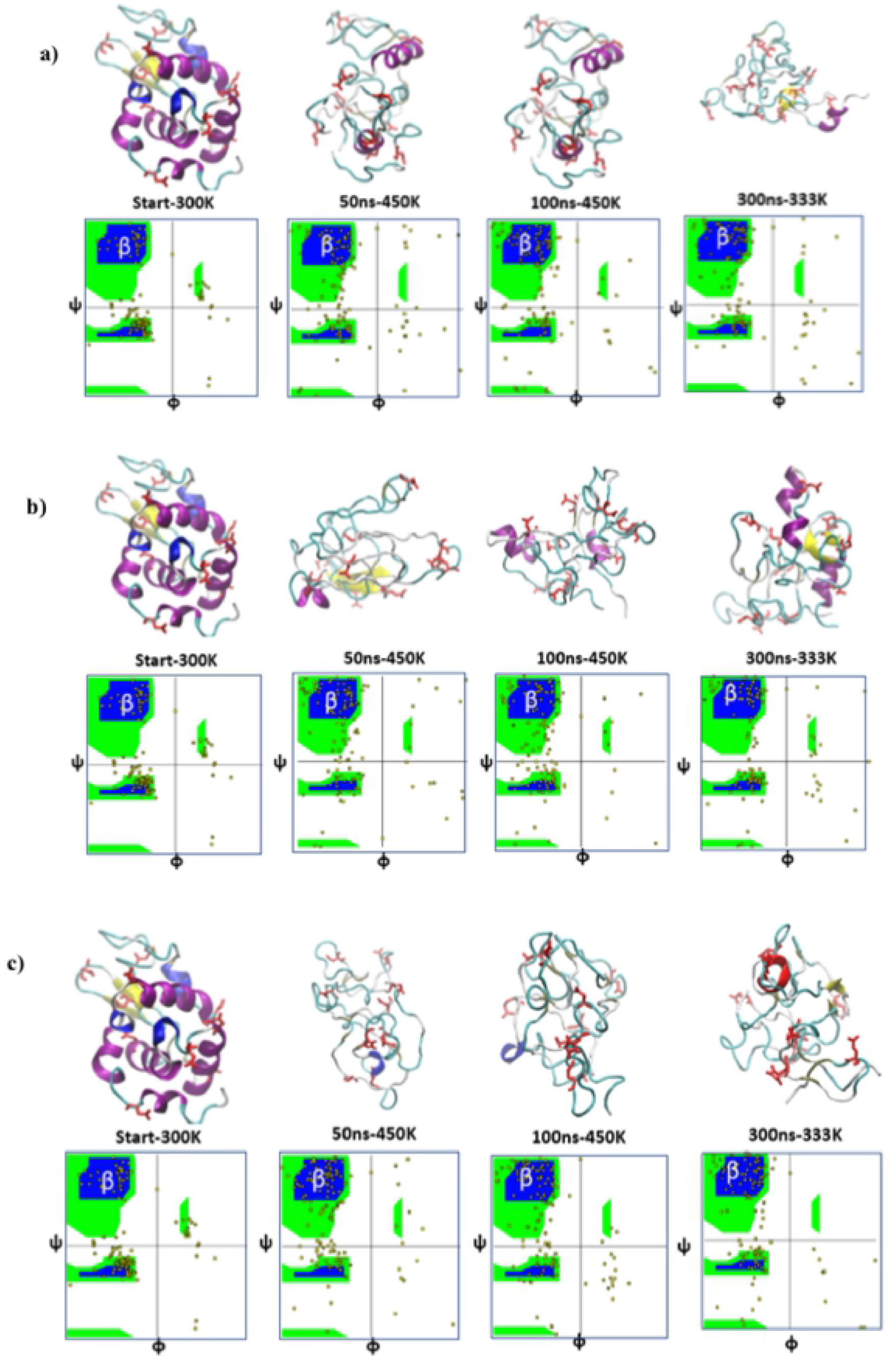
Conformational snapshots of HEWL at pH7 (top) and ramachandran plots (bottom) for the replicas (a) R0, (b) R1, and (c) R2 taken from at the start, after 50 ns of simulations at 450 K, after 100 ns of simulations at 450 K, and after 300 ns of simulations at 333 K.

The transitions from alpha-helixes, which were the major part of native lysozymes, into beta-strands and random coils were quantified through the Ramachandran plot. In the plots, backbone dihedral angles *φ* and *ψ* for each amino acid residue were mapped for the conformational snapshots at the start of the simulations, after 50 ns of simulations at 450 K, after 100 ns of simulations at 450 K, and the final snapshots after 300 ns of simulations at 333 K. In Figure 2 and Figure 3, *φ* and *ψ* angles of Ramachandran plots were displayed within the ranges between -180° to +180°. The regions within (*φ, ψ*) space of alpha-helical structures were around *φ* = -57° and *ψ* = -47° [38], while the regions for beta-strands were around *φ* = - 130° and *ψ* = +140°. For the three HEWL simulations at pH 2, the states of some amino acid residues were shifted from the alpha-helix region within the phase space to the beta-strand region, while some other amino acids were found shifted to other regions that corresponded to either the random-coils or the intermediate states between the alpha-helices and the beta-strands. The alpha-beta transitions occurred less frequently for the non-protonated proteins at pH7 (Figure 3), in which a large number of amino acid residues found at the intermediate states signified the incomplete refolding process as observed from the conformational snapshots.

The conformational changes were also monitored by the DSSP (Define Secondary Structure of Protein) algorithms (Figure 4 and 5) [37], displaying the plots of different secondary structure contents as a function of amino acid position (residue) and time. When the atomistic coordinates from x-ray crystallographic data or MD simulations are provided, DSSP can identify the types of secondary structures for all the amino acid residues from patterns of hydrogen bonding network. DSSP provided more quantitative analysis on the alpha-beta transition observed in our simulations at different pH. The analysis on the starting structure showed that the alpha-helix content within the native HEWL was 32.5% of the whole structure at pH 2, and 34.1% at pH 7 (Figure 5). Meanwhile, the beta-sheet or beta-strand content within the native HEWL was 6.2% of the whole structure at both pH values. During 300-ns simulations at 450 K and 333 K, the structures of HEWL were unfolded and then refolded, causing the percentage of alpha-helix and beta-sheet contents to change for all replicas. After the simulations finished, the percentage of alpha-helix content was found between 0.0% - 4.4% at pH2 and 0.1% - 13.0% at pH 7, suggesting that the full protonation on HEWL at pH 2 (Figure 4) corresponded to more alpha helix loss. However, the percentage of beta-sheet content was found between 9.1% - 19.1% at pH 2 and 0.6% - 3.8% at pH 7, suggesting that alpha-beta transition was more likely to occur at lower pH.

**Figure 4.**
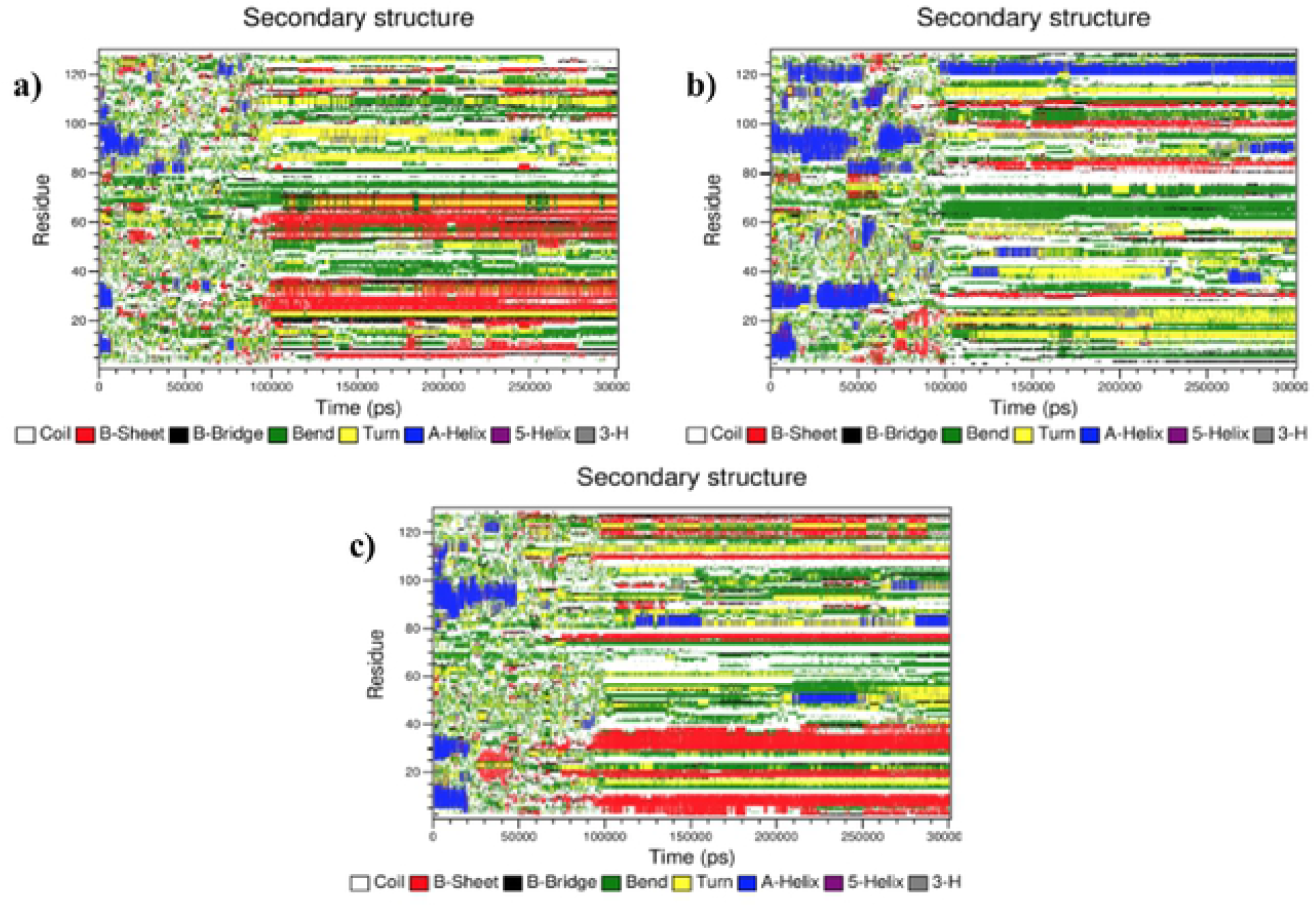
Time evolution of the secondary structures through the DSSP algorithm for the replica (a) R0, (b) R1, and (c) R2 at pH 2

**Figure 5.**
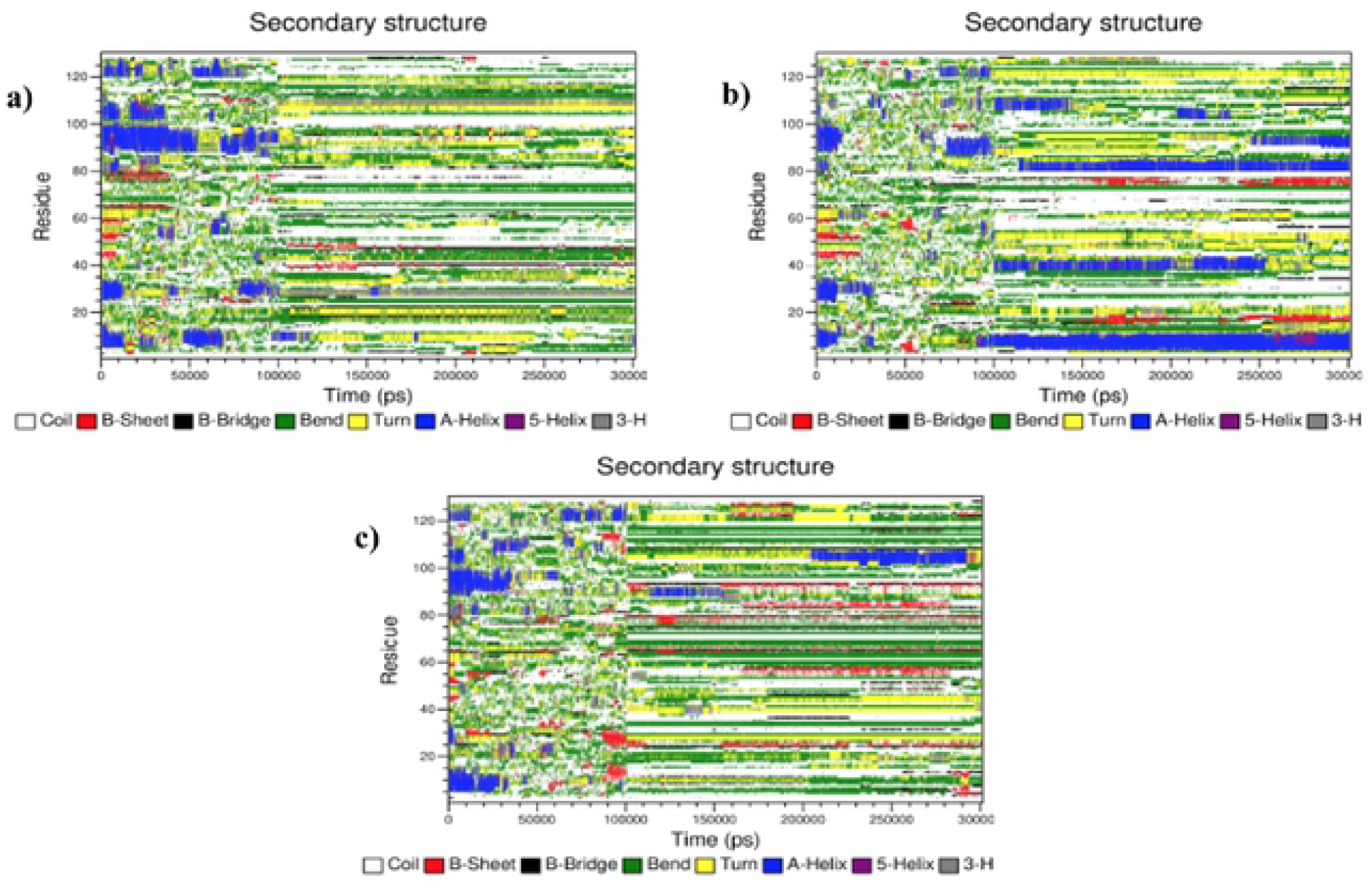
Time evolution of the secondary structures through the DSSP algorithm for the replica (a) R0, (b) R1, and (c) R2 at pH 7

In summary, our analysis in Figure 2 to Figure 5 was to monitor the conformational changes of HEWL during the MD simulations at pH 2 and pH 7 to observed the effects of protonation at low pH on the unfolding, refolding and alpha-beta transition for the propensity of amyloidosis. Conformational snapshots were taken, and Ramachandran plots were created at 50 ns and 100 ns under an elevated temperature of 450 K to observe the unfolding stages and at 300 ns under the lower temperature of 333 K to observe the refolding into different secondary structures. Beta-strands were more likely to form at pH 2, while at pH 7 the protein tended to either preserve alpha helices or unfold into random coils.

To elucidate the mechanism of beta-strand formation at pH2, the radius of gyration (Rg) was calculated as a function of time to monitor changes in the distribution of different compositions of HEWL, including hydrophobic, positively-charged, and negatively-charged amino acids affected by the protonations. Calculations of Rg were performed for the whole enzyme molecule and were performed separately for groups of positively-charged residues, negatively-charged residues, and hydrophobic residues from each simulation repeat at both 450 K (Figure 6) and 333 K (Figure 7) under both pH conditions. For the starting structure of HEWL at 450 K prior to unfolding, Rg of the positively charged group, the negatively charged group, and the hydrophobic group were different. Rg of the hydrophobic group was the lowest when compared with the other group as the hydrophobic sidechains were hidden within the inner core of the globular HEWL due to the hydrophobic effect, while Rg of the positively charged group was the highest Rg due to a large number of positive charge residues. The presence of positively-charged residues at the outer shell of HEWL, according to the highest Rg, was caused by Coulomb repulsions.

**Figure 6.**
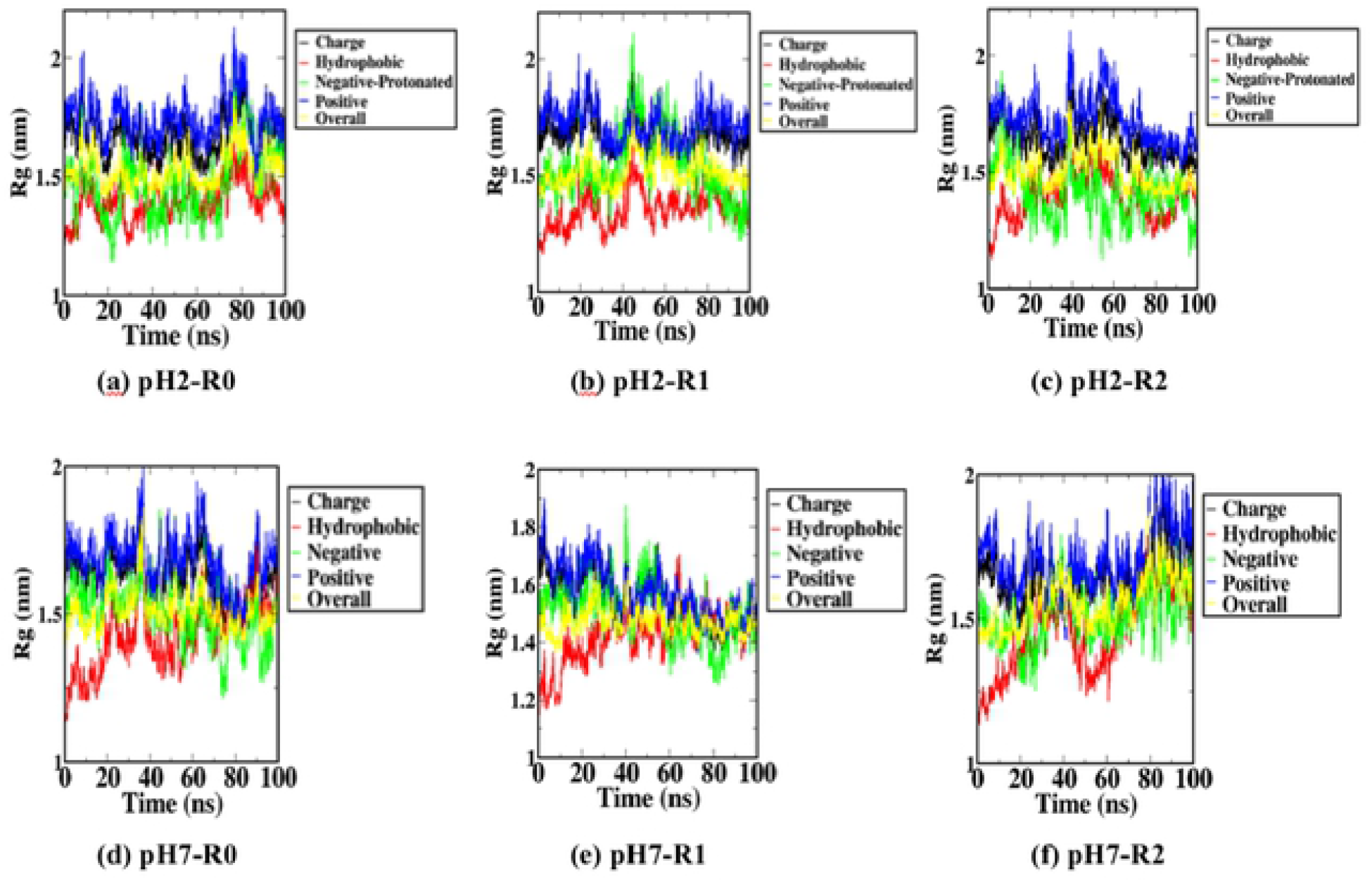
Radius of gyration calculated as functions of time for all three replaicas of HEWL at pH 2 (a, b and c) and pH 7 (d, e and t) at 450 K.

**Figure 7.**
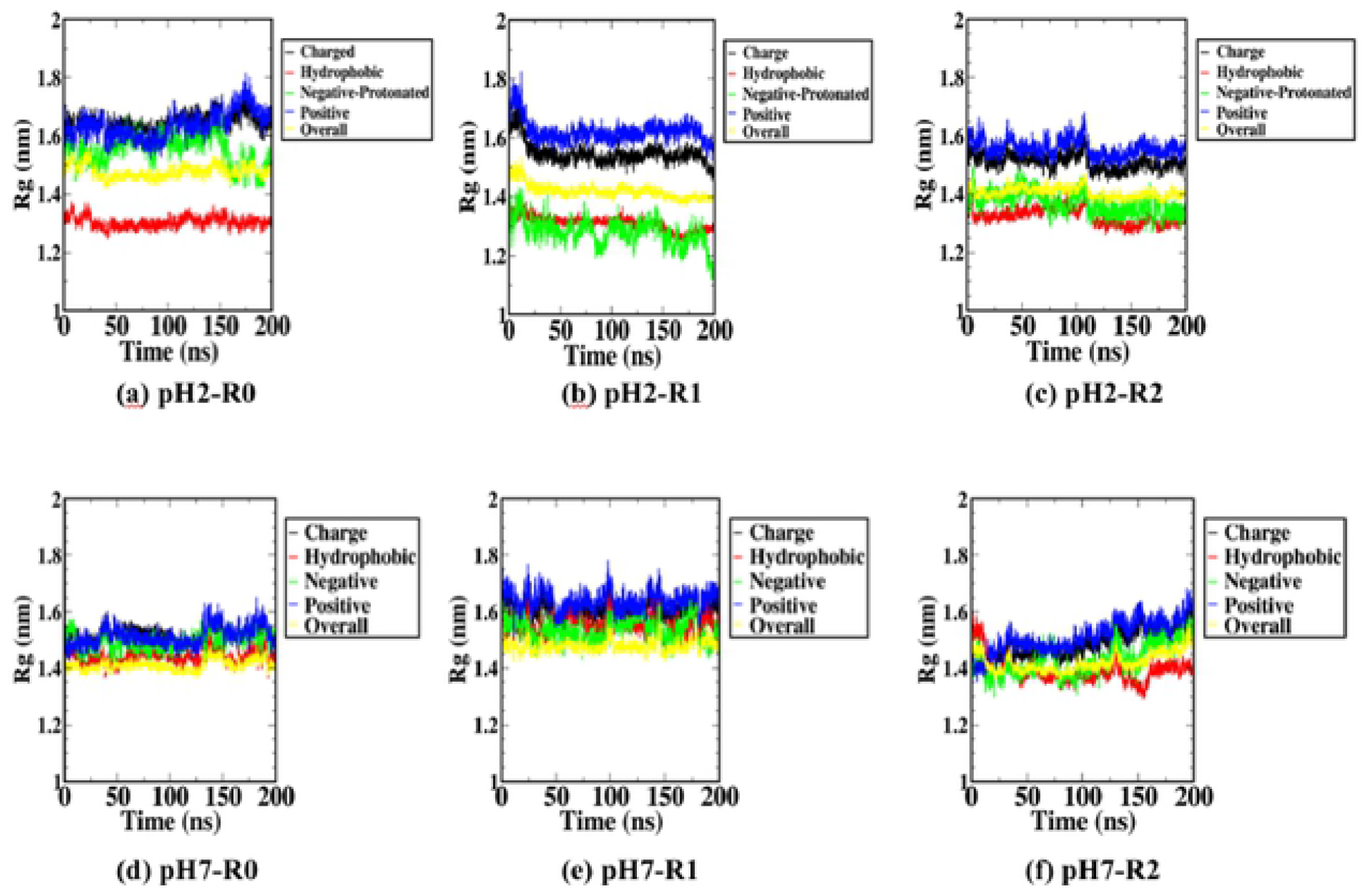
Radius of gyration calculated as functions of time for all three replaicas of HEWL at pH 2 (a, b and c) and pH 7 (d, e and f) at 333 K.

For all simulations at 450 K where unfolding occurred, the Rg profiles were highly fluctuated at both pH 2 and pH 7 due to the high level of thermal energy that disrupted the hydrogen bonds and overcame other molecular interactions. The distributions of positively-charged, negatively-charged, and hydrophobic compositions of HEWL at pH 2 (Figure 6a, 6b and 6c) could be differentiated by Rg values, similar to the native structure. However, at pH 7 (Figure 6d, 6e and 6f), Rg values of different amino acid groups tended to converge as the simulation progressed. The separation of Rg profile at pH 2 could be explained as the higher amount of positive charge caused stronger Coulomb repulsion between all positively charged residues. On the other hand, the mixing of Rg profiles between all amino acid groups at pH 7 suggested that the compositions of HEWL became more uniformly distributed as the protein unfolded. When the temperature of all simulations was reduced to 333 K, the Rg profiles became less fluctuated, but similar trends were preserved for both pH 2 (Figure 7a, 7b and 7c) and pH 7 (Figure 7d, 7e and 7f).

From Figure 8, clustering of the hydrophobic sidechains was observed underneath the beta-sheets formed at the surface of the refolded proteins at 333 K. The formation of hydrophobic clusters within the core of the protonated HEWL, as the protonation neutralized the negatively charged residues and caused the total charge of the system to be further positive. Coulomb repulsion between positively charged residues was the cause of larger Rg for the positively-charged amino acid group, corresponding to the presence of positively charged sidechains at the surface. As a result, the absence of positively charged residues from the core left an amount of space for hydrophobic sidechains to form hydrophobic clusters and thus lessen the Rg of the hydrophobic amino acid group. Meanwhile, the formation of beta-strands or beta-sheets from backbone parts of hydrophobic amino acids was facilitated by the increased compactness (low Rg) of the hydrophobic clusters.

**Figure 8.**
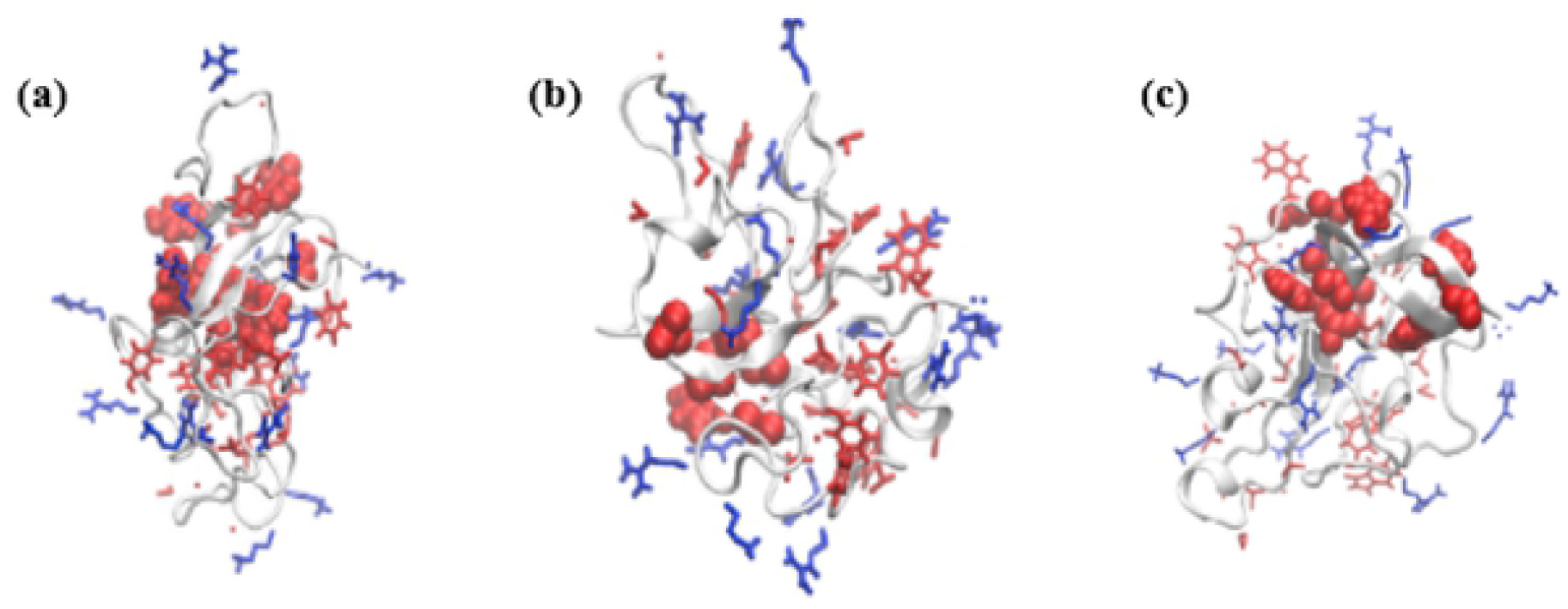
Beta strand formation of HEWL for all 3 replicas (a) R0, (b) R1, and (c) R2 under the pH 2 condition at 333 K. Hydrophobic sidechains were represented in red and positively charged sidechains were represented in blue, hydrophobic clusters were sho,vn by the van der Waals atomistic representation.

## 4. Discussions

A series of atomistic molecular dynamics (MD) simulations on HEWL under different pH conditions and conformational analysis of all simulation replicas were performed to understand the effect of low pH on the unfolding of HEWL and the propensity of refolding into the beta structure. For each replica, HEWL was explicitly simulated under the temperature 300 K, followed by 450 K and 333 K, in order to accelerate the unfolding and refolding processes to occur within a feasible timescale for MD simulations. RMSD was calculated for the simulations at both pH 2 and pH 7 to monitor the global conformational change or the unfolding of HEWL. No significant change was observed during the first period of all simulations at 300 K, while lysozymes at a high temperature of 450 K started to unfold from the beginning of all simulations under both pH conditions. Lysozymes at pH 2 unfolded faster compared to lysozymes at pH 7 due to the added positive charges from the amino acid protonations at low pH. When the temperature was decreased to 333 K, lysozymes tended to refold into beta-strands at pH 2, while misfolded into alpha-helices and random coils at pH 7. The transformation of alpha-helical structures of native lysozymes into beta-strands was also monitored by the Ramachandran plots. At pH 2, a large number of amino acid residues were moved from the alpha-helix region to the beta-strand region within the phase space of backbone angles, suggesting a large number of alpha-beta transitions. At pH 7, however, most of the amino acid residues of lysozyme tended to move to either the random-coil regions or to the intermediate states between the alpha-helix and the beta-strand regions. A similar trend was displayed from the time-dependent changes of the percentage of secondary structure contents quantified by the DSSP (Define Secondary Structure of Protein) algorithm. The analysis showed that the percentage of alpha-helix significantly decreased when lysozymes were unfolded at high temperatures for both pH conditions. However, at the final stages of the simulations, a higher amount of alpha-helix loss and a higher percentage of beta-strand formation were found at low pH. To elucidate the mechanism of these effects of low pH condition in terms of atomistic structure, the radius of gyration (Rg) was calculated for each of different compositions of HEWL, including the groups of hydrophobic, positively-charged, and negatively-charged amino acids. For the native HEWL structure, Rg values for the hydrophobic and the hydrophilic charged amino acid groups were different. Larger values of Rg were found for the charged amino acid groups, as the more hydrophilic charged amino acid groups energetically prefer the larger amount of contacts with water molecules and to stay further from the center of the protein. Meanwhile, smaller values of Rg were found for the hydrophobic amino acid groups, as they entropically prefer the lower amount of contacts with water molecules and to cluster closer to the center of the protein. For simulations at 450 K and pH 7, a substantial amount of kinetic energy was introduced with the elevated temperature. The kinetic energy from the global molecular motion overcame the interactions between the charged amino acids and the solvent molecules and the interactions between the hydrophobic sidechains. The random motion that occurred after the interaction network loss resulted in mixing between the hydrophilic and hydrophobic shells during the protein unfolding at pH 7. Interestingly, at pH 2, compositions of HEWL remained separated as hydrophobic and hydrophilic shells even at high temperatures. The separation was caused by the higher amount of Coulombic repulsion between 16 positively charged residues within the lysozymes. At Ph 7, the electrostatic repulsion was partly screened by the presence of the negatively charged amino acids, but this screening effect became vanished when the negatively charged amino acids were protonated at pH 2. As a result, positively charged residues formed a hydrophilic shell with larger Rg than the averaged Rg, and the hydrophobic residues stayed closed to the center of the protein globular structure due to the hydrophobic effect and possessed the lower Rg. The formation of beta-strands at the pH 2 condition can then be explained by the clustering of hydrophobic sidechains within the inner core. The higher amount of Coulombic repulsion at lower pH had driven most of the positively charged sidechain further from the backbone, leaving the backbone to stay at the middle between hydrophobic and hydrophilic shells. The beta-strands were finally formed by nucleation of the ordered backbone part.

## 5. Conclusions

In this study, the beta-strand formation mechanism at the early stage of amyloid fibrilization has been proposed. Lysozyme is one of the amyloid proteins that can misfold into fibrils and cause some diseases. However, the controlled synthetic amyloid fibrils can become useful biomaterials for bioengineering and biosensing applications. Our molecular dynamics simulations showed that beta-strands were more likely to form when HEWL was unfolded at pH 2. The mechanism of beta-strand formation was explained in terms of the radial distribution of the charged and the hydrophobic amino acids. Protonation of all glutamic acid and aspartic acid residues facilitates a higher amount of Coulomb repulsions. Positively charged side chains were separated from the cluster of hydrophobic side chains, and the backbone part formed beta-strands. Further validation is needed for the beta-strand formation of other amyloid proteins towards the control of fibril production and the prediction of nucleation sites.

## References

1. Xing L, Fan W, Chen N, Li M, Zhou X, Liu S. Amyloid formation kinetics of hen egg white lysozyme under heat and acidic conditions revealed by Raman spectroscopy. J Raman Spectrosc. 2019;50: 629–640. doi:10.1002/jrs.5567

2. Mishra R, Sörgjerd K, Nyström S, Nordigården A, Yu Y-C, Hammarström P. Lysozyme Amyloidogenesis Is Accelerated by Specific Nicking and Fragmentation but Decelerated by Intact Protein Binding and Conversion. J Mol Biol. 2007;366: 1029– 1044. doi:10.1016/j.jmb.2006.11.084

3. Yoshimura Y, Lin Y, Yagi H, Lee Y-H, Kitayama H, Sakurai K, et al. Distinguishing crystal-like amyloid fibrils and glass-like amorphous aggregates from their kinetics of formation. Proc Natl Acad Sci. 2012;109: 14446–14451. doi:10.1073/pnas.1208228109

4. Sabate R. When amyloids become prions. Prion. 2014;8: 233–239. doi:10.4161/19336896.2014.968464

5. Chiti F, Dobson CM. Protein Misfolding, Functional Amyloid, and Human Disease. Annu Rev Biochem. 2006;75: 333–366. doi:10.1146/annurev.biochem.75.101304.123901

6. Deidda G, Jonnalagadda SVR, Spies JW, Ranella A, Mossou E, Forsyth VT, et al. Self-Assembled Amyloid Peptides with Arg-Gly-Asp (RGD) Motifs As Scaffolds for Tissue Engineering. ACS Biomater Sci Eng. 2017;3: 1404–1416. doi:10.1021/acsbiomaterials.6b00570

7. Das S, Jacob RS, Patel K, Singh N, Maji SK. Amyloid Fibrils: Versatile Biomaterials for Cell Adhesion and Tissue Engineering Applications. Biomacromolecules. 2018;19: 1826–1839. doi:10.1021/acs.biomac.8b00279

8. Hauser CAE, Maurer-Stroh S, Martins IC. Amyloid-based nanosensors and nanodevices. Chem Soc Rev. 2014;43: 5326. doi:10.1039/C4CS00082J

9. Peralta MDR, Karsai A, Ngo A, Sierra C, Fong KT, Hayre NR, et al. Engineering Amyloid Fibrils from β-Solenoid Proteins for Biomaterials Applications. ACS Nano. 2015;9: 449–463. doi:10.1021/nn5056089

10. Altunbas A, Pochan DJ. Peptide-Based and Polypeptide-Based Hydrogels for Drug Delivery and Tissue Engineering. In: Deming T, editor. Peptide-Based Materials. Berlin, Heidelberg: Springer Berlin Heidelberg; 2011. pp. 135–167. doi:10.1007/128_2011_206

11. Jacob RS, Ghosh D, Singh PK, Basu SK, Jha NN, Das S, et al. Self healing hydrogels composed of amyloid nano fibrils for cell culture and stem cell differentiation. Biomaterials. 2015;54: 97–105. doi:10.1016/j.biomaterials.2015.03.002

12. Li C, Qin R, Liu R, Miao S, Yang P. Functional amyloid materials at surfaces/interfaces. Biomater Sci. 2018;6: 462–472. doi:10.1039/C7BM01124E

13. Seker UOS, Chen AY, Citorik RJ, Lu TK. Synthetic Biogenesis of Bacterial Amyloid Nanomaterials with Tunable Inorganic–Organic Interfaces and Electrical Conductivity. ACS Synth Biol. 2017;6: 266–275. doi:10.1021/acssynbio.6b00166

14. Kim J-Y, Sahu S, Yau Y-H, Wang X, Shochat SG, Nielsen PH, et al. Detection of Pathogenic Biofilms with Bacterial Amyloid Targeting Fluorescent Probe, CDy11. J Am Chem Soc. 2016;138: 402–407. doi:10.1021/jacs.5b11357

15. Humenik M, Scheibel T. Nanomaterial Building Blocks Based on Spider Silk– Oligonucleotide Conjugates. ACS Nano. 2014;8: 1342–1349. doi:10.1021/nn404916f

16. Bleem A, Daggett V. Structural and functional diversity among amyloid proteins: Agents of disease, building blocks of biology, and implications for molecular engineering: Amyloid Proteins: Beyond Disease. Biotechnol Bioeng. 2017;114: 7–20. doi:10.1002/bit.26059

17. Mankar S, Anoop A, Sen S, Maji SK. Nanomaterials: amyloids reflect their brighter side. Nano Rev. 2011;2: 6032. doi:10.3402/nano.v2i0.6032

18. Defelice FG, Ferreira ST. Physiopathological modulators of amyloid aggregation and novel pharmacological approaches in Alzheimer’s disease. An Acad Bras Ciênc. 2002;74: 265–284. doi:10.1590/S0001-37652002000200006

19. Chen Z, Li L, Zhao H, Guo L, Mu X. Electrochemical impedance spectroscopy detection of lysozyme based on electrodeposited gold nanoparticles. Talanta. 2011;83: 1501–1506. doi:10.1016/j.talanta.2010.11.042

20. Bogomolova A, Komarova E, Reber K, Gerasimov T, Yavuz O, Bhatt S, et al. Challenges of Electrochemical Impedance Spectroscopy in Protein Biosensing. Anal Chem. 2009;81: 3944–3949. doi:10.1021/ac9002358

21. Randviir EP, Banks CE. Electrochemical impedance spectroscopy: an overview of bioanalytical applications. Anal Methods. 2013;5: 1098. doi:10.1039/c3ay26476a

22. Knowles TPJ, Mezzenga R. Amyloid Fibrils as Building Blocks for Natural and Artificial Functional Materials. Adv Mater. 2016;28: 6546–6561. doi:10.1002/adma.201505961

23. Wang W. Protein aggregation and its inhibition in biopharmaceutics. Int J Pharm. 2005;289: 1–30. doi:10.1016/j.ijpharm.2004.11.014

24. Sutthibutpong T, Rattanarojpong T, Khunrae P. Effects of helix and fingertip mutations on the thermostability of xyn11A investigated by molecular dynamics simulations and enzyme activity assays. J Biomol Struct Dyn. 2018;36: 3978–3992. doi:10.1080/07391102.2017.1404934

25. Moraitakis G, Goodfellow JM. Simulations of Human Lysozyme: Probing the Conformations Triggering Amyloidosis. Biophys J. 2003;84: 2149–2158. doi:10.1016/S0006-3495(03)75021-8

26. Robustelli P, Piana S, Shaw DE. Developing a molecular dynamics force field for both folded and disordered protein states. Proc Natl Acad Sci. 2018;115: E4758–E4766. doi:10.1073/pnas.1800690115

27. Guvench O, MacKerell AD. Comparison of Protein Force Fields for Molecular Dynamics Simulations. In: Kukol A, editor. Molecular Modeling of Proteins. Totowa, NJ: Humana Press; 2008. pp. 63–88. doi:10.1007/978-1-59745-177-2_4

28. Jafari M, Mehrnejad F. Molecular Insight into Human Lysozyme and Its Ability to Form Amyloid Fibrils in High Concentrations of Sodium Dodecyl Sulfate: A View from Molecular Dynamics Simulations. Hassan I, editor. PLOS ONE. 2016;11: e0165213. doi:10.1371/journal.pone.0165213

29. Jiang Z, You L, Dou W, Sun T, Xu P. Effects of an Electric Field on the Conformational Transition of the Protein: A Molecular Dynamics Simulation Study. Polymers. 2019;11: 282. doi:10.3390/polym11020282

30. Walker AR, Baddam N, Cisneros GA. Unfolding Pathways of Hen Egg-White Lysozyme in Ethanol. J Phys Chem B. 2019;123: 3267–3271. doi:10.1021/acs.jpcb.9b01694

31. Sedov IA, Magsumov TI. Molecular dynamics study of unfolding of lysozyme in water and its mixtures with dimethyl sulfoxide. J Mol Graph Model. 2017;76: 466–474. doi:10.1016/j.jmgm.2017.07.032

32. Patel D, Kuyucak S. Computational study of aggregation mechanism in human lysozyme[D67H]. Zhang Y, editor. PLOS ONE. 2017;12: e0176886. doi:10.1371/journal.pone.0176886

33. Liu H-L, Wu Y-C, Zhao J-H, Fang H-W, Ho Y. Structural Analysis of Human Lysozyme Using Molecular Dynamics Simulations. J Biomol Struct Dyn. 2006;24: 229–238. doi:10.1080/07391102.2006.10507115

34. English NJ, Mooney DA. Denaturation of hen egg white lysozyme in electromagnetic fields: A molecular dynamics study. J Chem Phys. 2007;126: 091105. doi:10.1063/1.2515315

35. Søndergaard CR, Olsson MHM, Rostkowski M, Jensen JH. Improved Treatment of Ligands and Coupling Effects in Empirical Calculation and Rationalization of p K a Values. J Chem Theory Comput. 2011;7: 2284–2295. doi:10.1021/ct200133y

36. Olsson MHM, Søndergaard CR, Rostkowski M, Jensen JH. PROPKA3: Consistent Treatment of Internal and Surface Residues in Empirical p K a Predictions. J Chem Theory Comput. 2011;7: 525–537. doi:10.1021/ct100578z

37. Kabsch W, Sander C. Dictionary of protein secondary structure: Pattern recognition of hydrogen-bonded and geometrical features. Biopolymers. 1983;22: 2577–2637. doi:10.1002/bip.360221211

38. Hovmöller S, Zhou T, Ohlson T. Conformations of amino acids in proteins. Acta Crystallogr D Biol Crystallogr. 2002;58: 768–776. doi:10.1107/S0907444902003359

